# Kinetics and identities of extracellular peptidases in subsurface sediments of the White Oak River Estuary, NC

**DOI:** 10.1101/080671

**Authors:** Andrew D. Steen, Richard T. Kevorkian, Jordan T. Bird, Nina Dombrowski, Brett J. Baker, Shane M. Hagen, Katherine H. Mulligan, Jenna M. Schmidt, Austen T. Webber, Taylor Royalty, Marc J. Alperin

## Abstract

Anoxic subsurface sediments contain communities of heterotrophic microorganisms that metabolize organic carbon at extraordinarily slow rates. In order to assess the mechanisms by which subsurface microorganisms access detrital sedimentary organic matter, we measured kinetics of a range of extracellular peptidases in anoxic sediments of the White Oak River estuary, NC. Nine distinct peptidase substrates were enzymatically hydrolyzed at all depths. Potential peptidase activities (*V_max_*) decreased with increasing sediment depth, although *V_max_* expressed on a per cell basis was approximately the same at all depths. Half-saturation constants (*K_m_*) decreased with depth, indicating peptidases that functioned more efficiently at low substrate concentrations. Potential activities of extracellular peptidases acting on molecules that are enriched in degraded organic matter (D-phenylalanine and L-ornithine) increased relative to enzymes that act on L-phenylalanine, further suggesting microbial community adaptation to access degraded organic matter. Nineteen classes of predicted, exported peptidases were identified in genomic data from the same site, of which genes for class C25 (gingipain-like) peptidases represented more than 40% at each depth. Methionine aminopeptidases, zinc carboxypeptidases, and class S24-like peptidases, which are involved in single-stranded DNA repair, were also abundant. These results suggest a subsurface heterotrophic microbial community that primarily accesses low-quality detrital organic matter via a diverse suite of well-adapted extracellular enzymes.

**Importance:** Burial of organic carbon in marine and estuarine sediments represents a long-term sink for atmospheric carbon dioxide. Globally, ∼40% of organic carbon burial occurs in anoxic estuaries and deltaic systems. However, the ultimate controls on the amount of organic matter that is buried in sediments, versus oxidized into CO_2_, are poorly constrained. Here we used a combination of enzyme assays and metagenomic analysis to identify how subsurface microbial communities catalyze the first step of proteinaceous organic carbon degradation. Our results show that microbial communities in deeper sediments are adapted to access molecules characteristic of degraded organic matter, suggesting that those heterotrophs are adapted to life in the subsurface.

## Introduction

A large fraction of the microorganisms in subsurface sediments are heterotrophs that metabolize aged, microbially altered organic matter (1–3). These communities’ metabolisms can be more than a million-fold more slower than cells in culture (1, 4). A recent meta-analysis showed that only about 12% of cells in marine sediments belonged to cultured species, while 27% belong to phyla that contain no cultured representatives (5). Consequently, the mechanisms by which these microorganisms access detrital organic matter are poorly understood (6).

In surface environments, where organic carbon metabolism is relatively rapid, heterotrophic microorganisms gain energy by metabolizing a combination of small molecules (<600-1000 Da), which can be taken up directly via general uptake porins (7) and macromolecules, which must be broken down outside of the cell by extracellular enzymes. Most freshly-produced organic matter is macromolecular, and large molecules tend to be more bioavailable than small ones (8), so the nature and activity of extracellular enzymes present in surface environments is a major control on the rate of microbial carbon oxidation in such environments.

It is not clear whether microbial extracellular enzymes play the same role in subsurface sediments. It is conceivable that macromolecules are broken down primarily by non-enzymatic mechanisms in sediments. For instance, in soils, MnO_4_ catalyzes the depolymerization of proteins without requiring enzymes. Certain bacterial species can use TonB-dependant transporters to transport polysaccharides that are substantially larger than 600 Da into the periplasm (although enzymatic hydrolysis is still required prior to uptke into the cytoplasm; 9). Furthermore, some of the unique aspects of subsurface sediments suggest that extracellular enzymes might not be an effective strategy to obtain carbon or energy. In order for the production of extracellular enzymes to be part of a viable metabolic strategy, each enzyme must, over its lifetime, provide the cell with at least as much carbon or energy as was required to synthesize the enzyme (10–12). In subsurface sediments, where cell division times may be on the order of decades to millenia, enzyme lifetimes would need to be correspondingly longer to remain ‘profitable’. Since enzyme lifetimes are finite, there must exist a community metabolic rate below which extracellular enzyme lifetimes are too short to become profitable. That limit is difficult to quantify because enzyme lifetimes in any environment are poorly constrained (e.g., 13). Thus, it is plausible that extracellular enzyme-mediated carbon acquisition is impractical in sediments in which metabolic rates are particularly slow.

While extracellular enzyme activity in surface sediments has frequently been reported, few reports exist of extracellular enzyme activity from deeper than 20 cm below the seafloor (cmbsf; 14). Enzyme activity has been reported in sapropels up to 389 cmbsf in sapropels in the Eastern Mediterranean Sea (15, 16) and in sediment from 600-630 cmbsf in Aarhus Bay sediments (17), as well as in a few other subsurface environments, such as the interior of seafloor basalts at the Loihi seamount (18). Furthermore, an analysis of transcriptomes from subsurface sediments of the Peru Margin revealed diverse exported peptidases and carbohydrate-active enzymes, which decreased in relative abundance with increasing depth (19).

In order to better understand how heterotrophic microorganisms in subsurface sediments access organic matter, we assayed a diverse set of peptidases (protein-degrading enzymes) in sediment cores from the White Oak River estuary, NC. We paired these assays with analysis of the potential for extracellular peptidase production from existing metagenomic data sets. We chose this site because the porewater geochemistry and microbiology of these sediments has been well-characterized (20–24) and because they contain abundant Bathyarchaeota and Thermoprofundales archaea, which appear to be capable of metabolizing detrital organic matter (17, 25–27). We focused on peptidases because protein degradation appears to be an important metabolism for some subsurface archaea (17) and because peptidases were more active than other enzymes in similar environments (15, 18). Because environmental samples contain a wide range of distinct peptidases at variable activities (28, 29) we measured the hydrolysis of eleven different substrates, which may be hydrolyzed by structurally and genetically diverse extracellular peptidases.

## Results

### Peptidase kinetics

Sediment cores were sampled on two dates: first, to measure saturation curves for six structurally diverse peptidase substrates, and second to measure a more targeted set of peptidases at high depth resolution in order to assess the relative ability of subsurface communities to access more degraded organic matter. Combining all samples, unambiguous hydrolysis of nine different peptidase substrates was observed. All peptidase substrates assayed with the more-sensitive single-cuvette methodology were hydrolyzed much faster in untreated sediments than in autoclaved controls (Fig 1). Kinetics of substrate hydrolysis were qualitatively consistent with the Michaelis-Menten rate law, 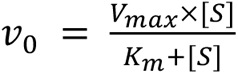, with estimated *V_max_* values ranging from 40 to 3400 nmol g^-1^ sediment hr^-1^ (median 310, interquartile range 190 to 560 nmol g^-1^ sediment hr^-1^). Throughout the core, AAF-AMC, GGR-AMC, and Gly-AMC were hydrolyzed the fastest, and Arg-AMC was hydrolyzed slowest (Fig 2). Summed *V_max_* values for each substrate, a proxy for the total peptidolytic potential of the microbial community, decreased with depth from 9.09 μmol AMC g^-1^ sed hr^-1^ at the surface to 1.24 μmol AMC g^-1^ sed hr^-1^, or 13% of the surface value, at 82.5 cmbsf. Estimated *K_m_* values ranged from 36.1 μM to 1310 μM (median 138 μM, interquartile range 102 to 326 μM), and trended downward (i.e., to greater substrate affinity) with increasing depth (Fig 3). *K_m_* values for hydrolysis of Leu-AMC were the greatest (i.e., lowest substrate affinity) while *K_m_* values for hydrolysis of BocVPR-AMC, GGR-AMC, and Arg-AMC were the least.

**Figure 1:**
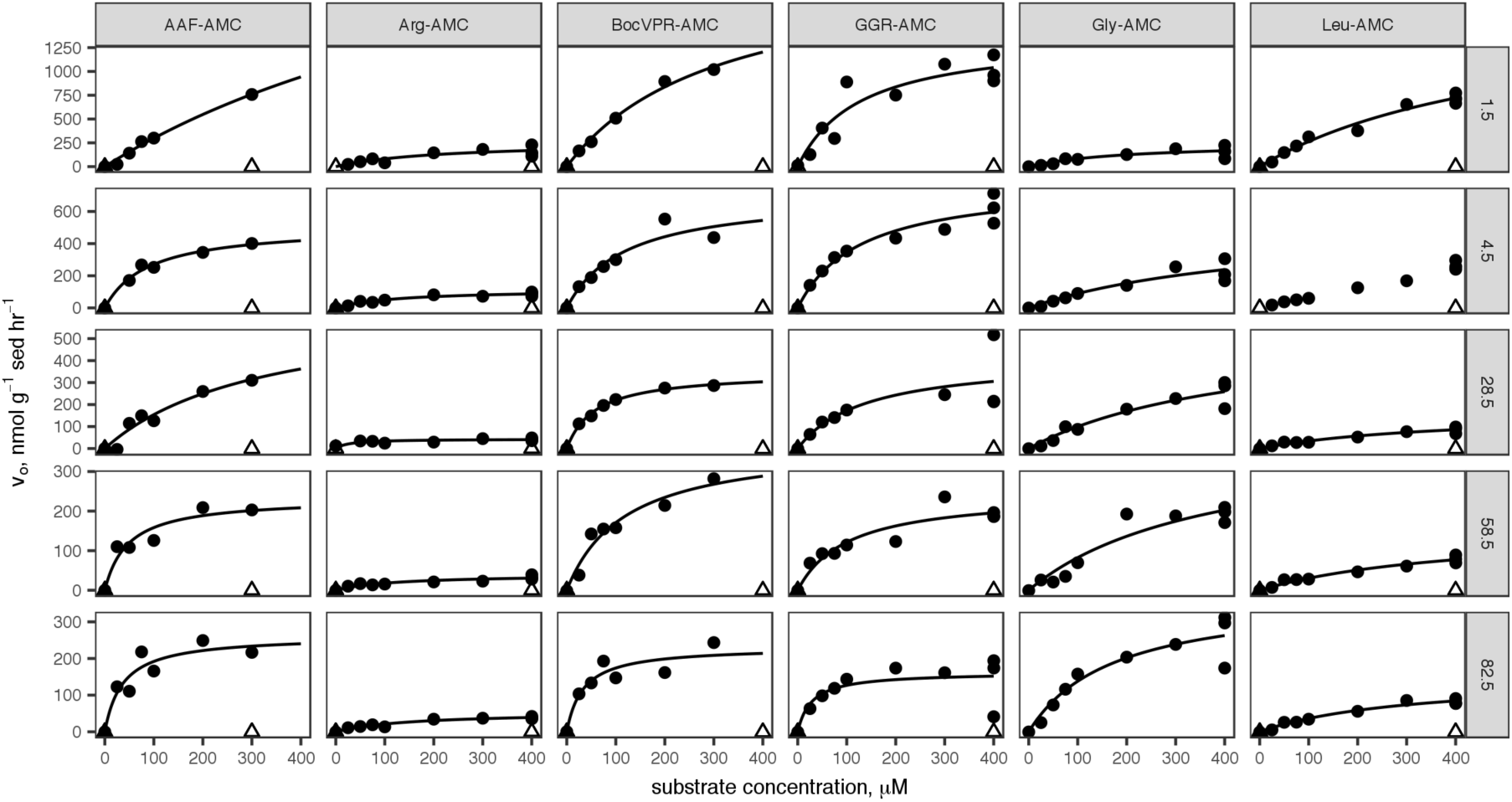
Saturation curves for six substrates measured using the single-cuvette reader methodology at each of six depths. Dark circles indicate “live” samples, open triangles indicate autoclaved controls. Lines indicate nonlinear least-squares fits to the Michaelis-Menten rate law. Substrate abbreviations are given in the column headings and are defined in Table 1. Sediment depths are listed on row headings in centimeters below sediment-water interface.

**Fig 2:**
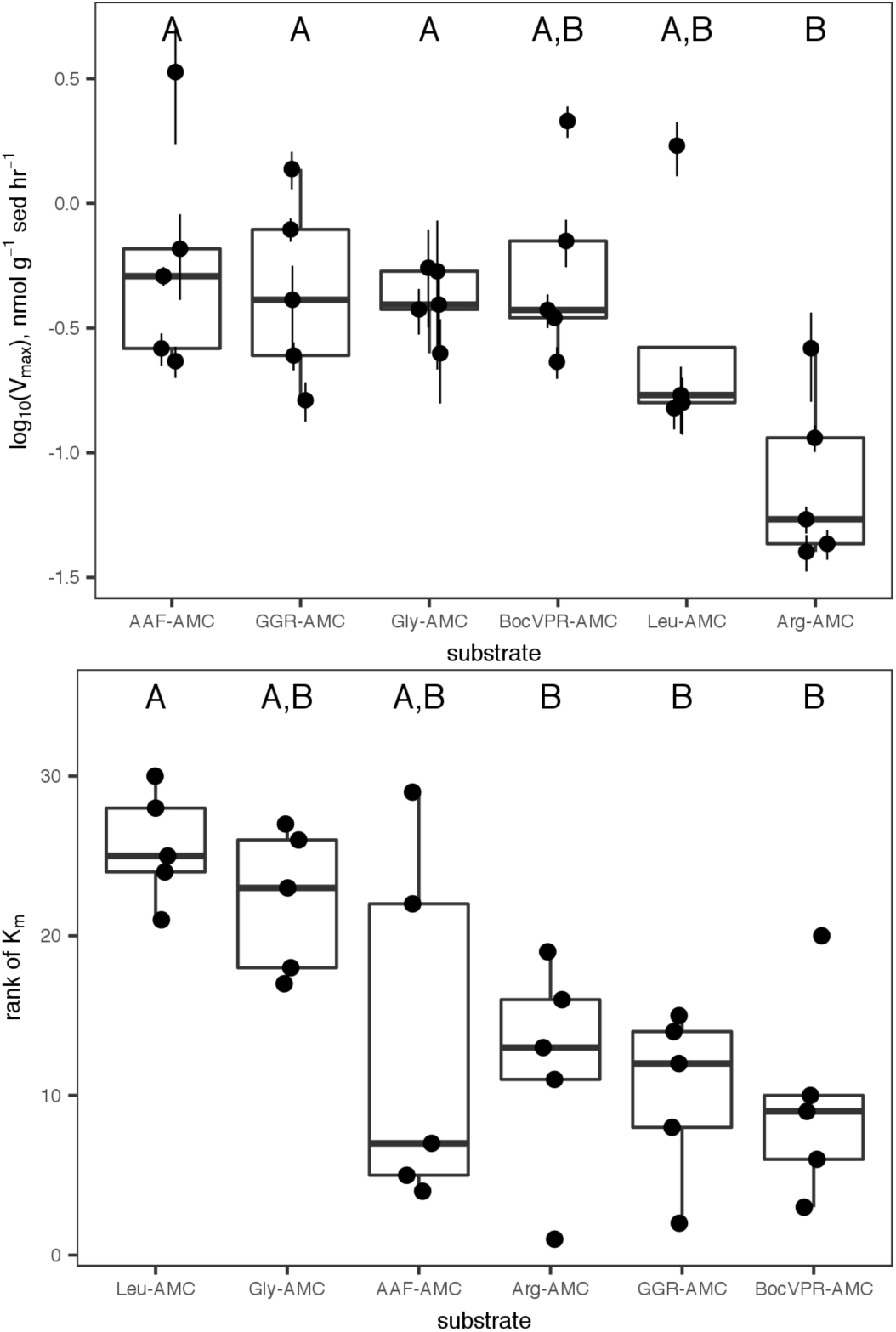
*V_max_* and *K_m_* values, shown individually with error bars indicating standard error of the nonlinear least-squares estimate, and collectively in a box-and-whiskers plot. Substrates sharing a letter are not significantly different according to one-way ANOVA of log_10_-transformed data with Tukey HSD post-hoc analysis (*V_max_*), or Kruskal-Wallis test with Tukey HSD post-hoc analysis (*K_m_*).

**Fig 3:**
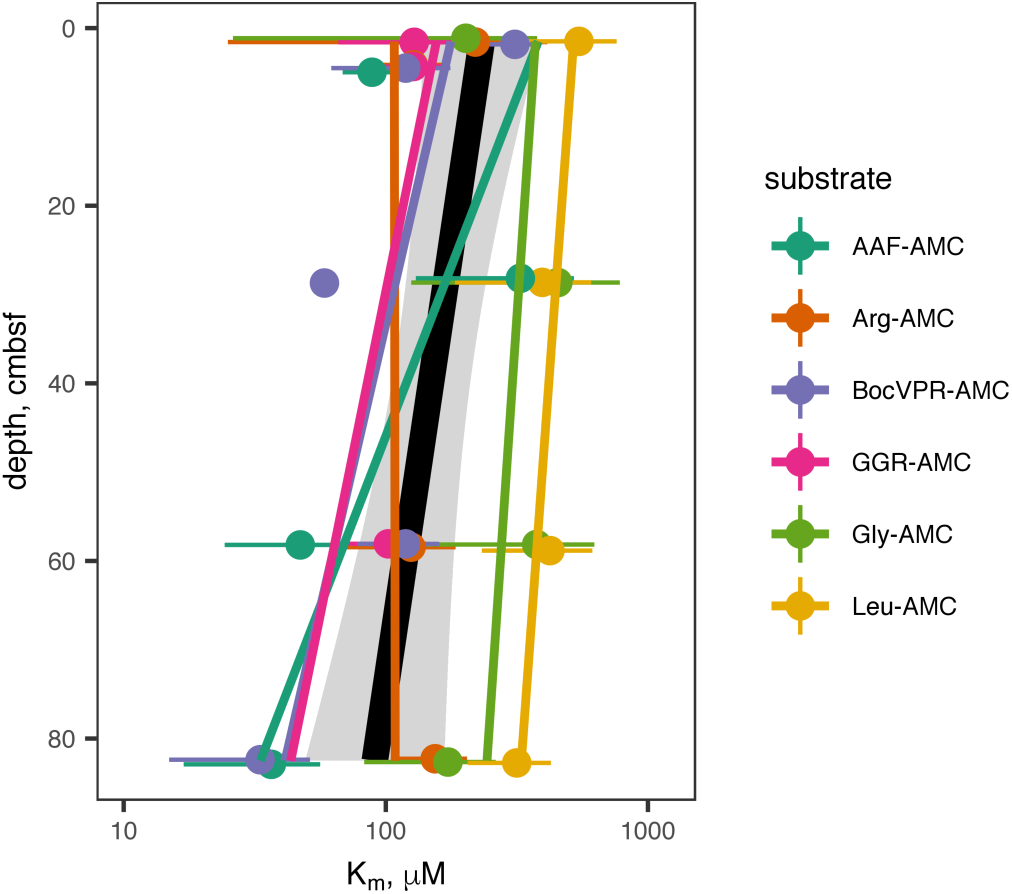
*K_m_* values of extracellular peptidases as a function of depth. Details of substrates and the peptidases they correspond to are in Table 1. Error bars represent the standard deviation of replicate samples. Colored lines represent a linear least-squares regression for each substrate. The black line and gray shading represent linear regression and 95% confidence interval for all substrates taken together.

**Table 1:** Substrates used in this study and the enzymes that hydrolyze them. AMC stands for aminomethylcoumarin, the moiety that becomes fluorescent after hydrolysis of the peptide bond. All amino acids are in the L-stereoconformation unless otherwise noted. Enzymes are described as “putative” because the substrate specificity of many environmental peptidases is fairly broad, so multiple peptidases may hydrolyze any given substrate.

| Substrate | Abbreviation | Putative enzyme |
| --- | --- | --- |
| L-arginine-7-amido-4-methylcoumarin | Arg-AMC | arginyl aminopeptidase |
| L-glycine-7-amido-4-methylcoumarin | Gly-AMC | glycyl aminopeptidase |
| L-leucine-7-amido-4-methylcoumarin | Leu-AMC | leucyl aminopeptidase |
| Carboxybenzoyl-glycine-glycine-arginine-7-amido-4-methylcoumarin | Z-GGR-AMC | gingipain and other endopeptidases |
| Alanine-alanine-phenylalanine-7-amido-4-methylcoumarine | AAF-AMC | clostripain and other endopeptidases |
| Boc-valine-proline-arginine-AMC | Boc-VPR-AMC | gingipain and other endopeptidases |
| D-phenylalanine-AMC | D-Phe-AMC | D-phenylalanine aminopeptidase |
| L-phenylalanine-AMC | L-Phe-AMC | L-phenylalanine-aminopeptidase |
| Ornithine-AMC | Orn-AMC | ornithine aminopeptidase |

In a separate core, hydrolysis rates of D-Phe-AMC, L-Phe-AMC, and L-Orn-AMC were assessed. These were measured using a plate-reader technique that proved insufficiently precise to accurately measure *V_max_* or *K_m_*, so we have only reported the observed hydrolysis rate *v_0_*, which was measured at a high substrate concentration (400 μM) and therefore approximates *V_max_*. Ratios of *v_0_* for D-Phe-AMC : L-Phe-AMC hydrolysis and L-Orn-AMC : L-Phe-AMC hydrolysis rates increased approximately linearly downcore (Fig 4).

**Fig 4:**
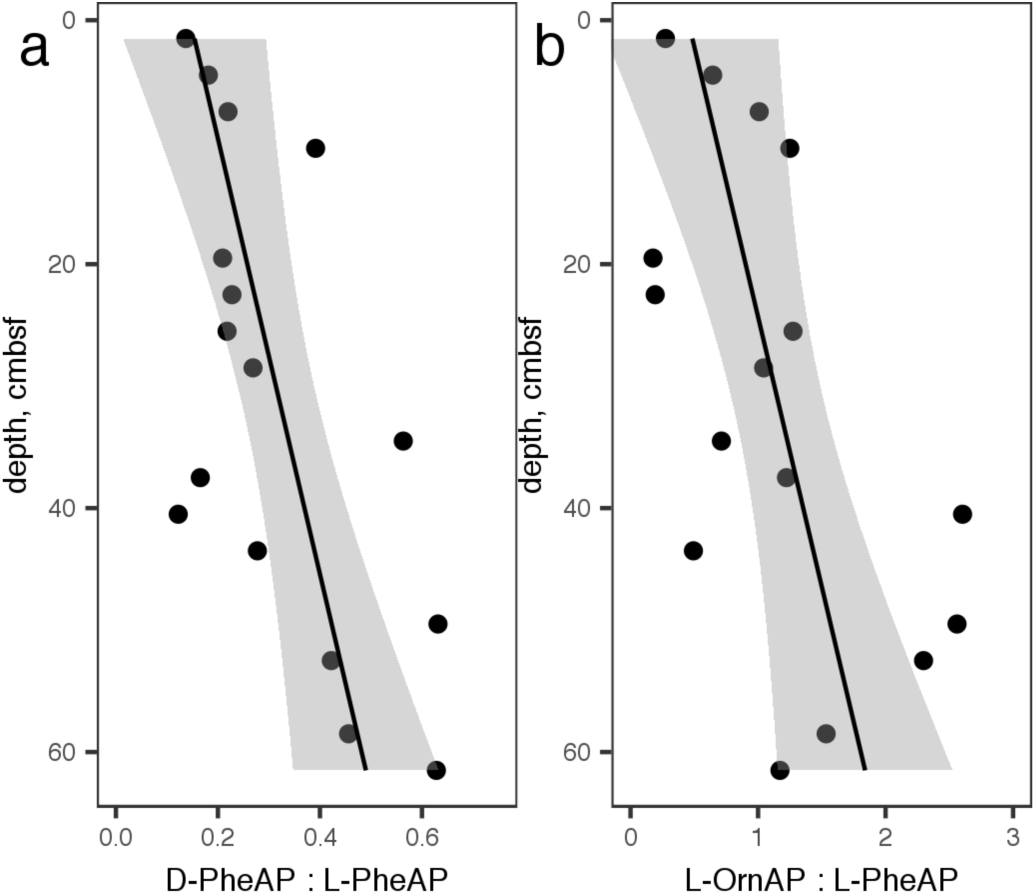
a.) Ratio of *v_0_* for D-phenylalanine aminopeptidase: L-phenylalanine aminopeptidase and b.) L-ornithine aminopeptidase : L-phenylalanine aminopeptidase. The linear regressions are given by D-Phe-AP:L-Phe-AP = 5.60 (± 1.87) × 10^-3^ × depth + 0.146 (± 0.067) and L-Orn-AP:L-Phe-AP = 2.26 (± 0.89) × 10^-3^ × depth + 0.451 (± 0.326)

### Microbial abundance, cell-specific peptidase activity, and organic carbon oxidation rates

Concordantly with potential activities (Fig 5a), cell counts decreased more-or-less steadily downcore from 4.5 × 10^8^ cells ml^-1^ wet sediment at 1.5 cmbsf to 7.4 × 10^7^ cells ml^-1^ wet sediment at 82.5 cmbsf. Consequently, cell-specific total potential peptidase activity was roughly constant at 32 +/-14 amol AMC cell^-1^ hr^-1^ (Fig 5b) with no significant trend as a function of depth. Most of the error in cell-specific peptidase activities results from variance in cell counts rather than in *V_max_* estimations.

**Figure 5:**
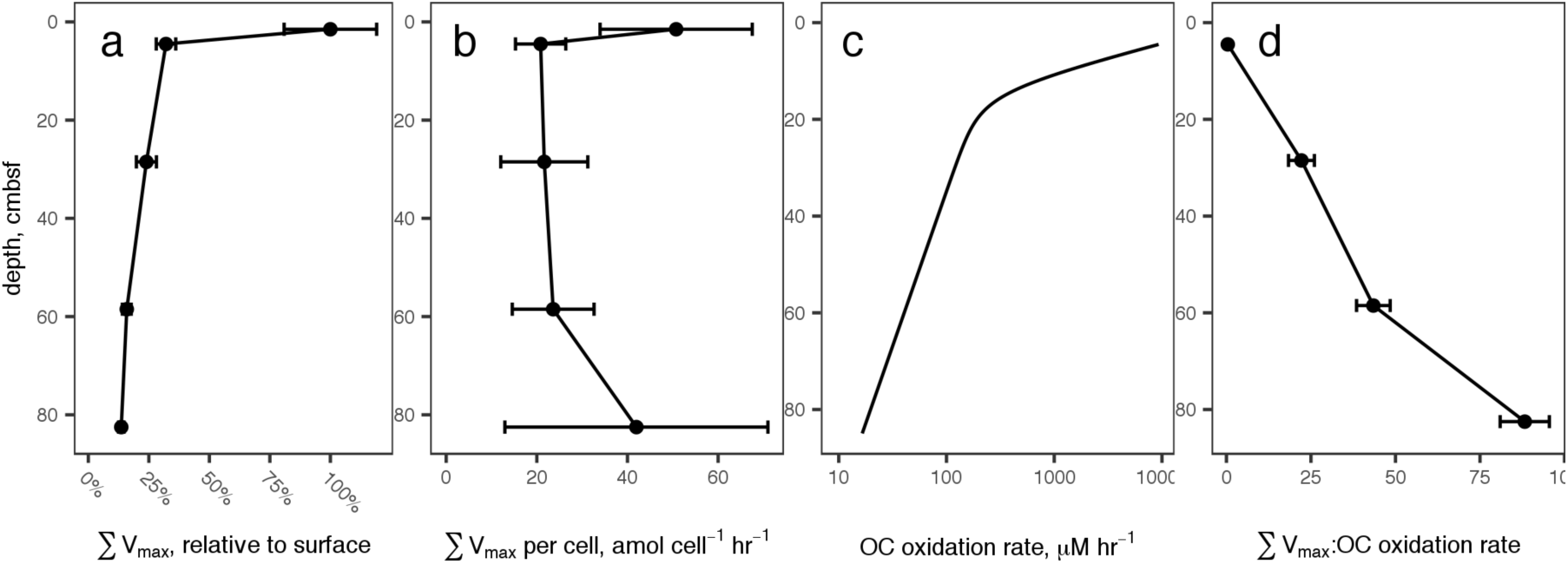
a.) The sum of all peptidase Vmax values, relative to the value at 4.5 cm, versus sediment depth. Error bars represent propagated error of the estimate of *V_max_* for each substrate. b.) Summed *V_max_* relative to cell count. Error bars represent propagated error from summed *V_max_* and cell counts, and is dominated by cell count uncertainty. c.) Organic carbon oxidation rates modeled from sulfate and methane profiles. d.) Summed *V_max_* relative to modeled carbon oxidation rates. Error bars represent error in summed *V_max_* relative to organic carbon oxidation rates, for which uncertainty was not modeled.

Organic carbon oxidation rates were estimated using a 2-G model driven by porewater methane and sulfate concentrations. The total modeled organic carbon oxidation rate *G* at 82.5 cmbsf was approximately 0.17% relative to that at 4.5 cmbsf (the top of the model domain), a decrease of almost 3 orders of magnitude (Fig 5c). Thus, summed *V_max_* relative to G, a proxy for the effort microbes exert compared to obtain complex organic carbon relative the amount of carbon they metabolize, increased more than 200-fold in the deepest sediments relative to surface sediments (Fig 5d).

### Peptidase genes and microbial taxa

Samples for genomic analysis were taken from three broad sedimentary zones: the sulfate reduction zone (SRZ, 8-12 cmbsf), sulfate-methane transition zone (SMTZ, two distinct samples from nearby locations, 24-32 cmbsf and 26-30 cmbsf), and the methane-rich zone (52-54 cmbsf; data originally published in Baker *et al.*, 2015). A total of 3739 genes encoding extracellular peptidases were identified among metagenomes from the three depth zones examined, including 685 from SRZ, 1994 from SMTZ, and 1060 from MRZ. Of the genes encoding for peptidases, 0-71% (depending on class of peptidase, algorithm and sediment depth) contained a signal peptide (SP) and are likely secreted by the SEC-dependent transport system (supplementary data). Among the genes associated with signal peptides, members of peptidase family C25, belonging to the gingipain family, were by far the most abundant at all depths, accounting for 41-45% of all SP-associated peptidases (Fig. 6a). Genes annotated as encoding extracellular methionine aminopeptidases and zinc carboxypeptidases were also abundant (13%-19%). Together, these peptidase classes accounted for 73%, 76%, and 73% of exported peptidases in the SRZ, SMTZ, and MRZ respectively. The composition of protein families was generally consistent with depth, particularly among the more-abundant peptidases. Five peptidase annotations were an exception to this trend: peptidase family M1, peptidase family M20/M25/M40, peptidase family M3, M61 glycyl aminopeptidase, and thermophilic metalloprotease (M29) were found in much lower abundance at the SMTZ than the MRZ or SRZ. Given that those correspond to differences of one or a few total reads, these are well within the range of noise.

**Fig 6:**
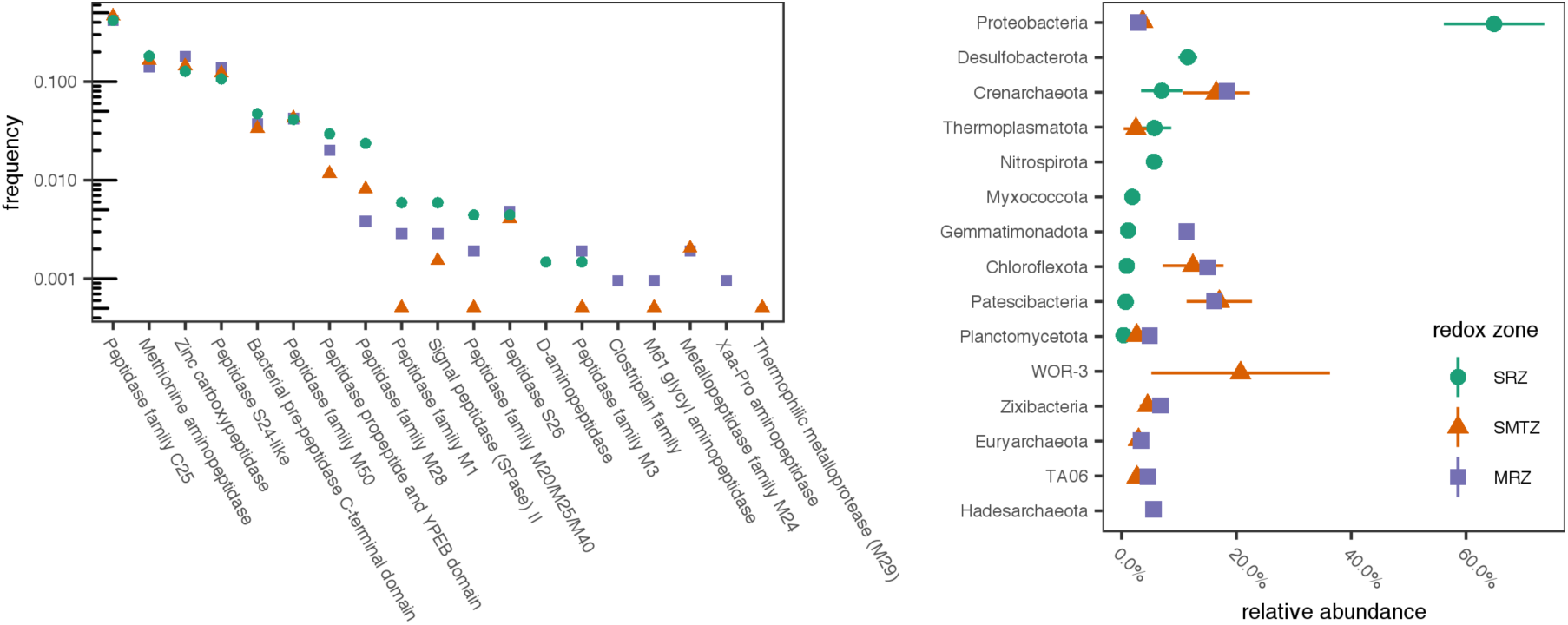
A.) Frequency of reads for genes of various classes of extracellular peptidases that were associated with signal peptidases, relative to all genes for extracellular peptidases at that depth. SRZ, SMTZ, and MRZ correspond to “sulfate reducing zone” (8-12 cmbsf), “sulfate-methane transition zone” (24-32 cmbsf) and “methane-rich zone” (24-28 cmbsf) respectively. B.) Relative abundance of phyla in bins at each depth. Only the 10 most abundant phyla at each depth are shown. The orange SMTZ points represent the average of two SMTZ samples, taken ∼500 m apart from each other, and error bars represent the range of the two sites.

*Proteobacteria* were by far the most abundant phylum in the SRZ sediments, with smaller contributions from *Chloroflexi*, *Bacteroidetes*, *Planctomycetes*, *Euryarchaeota*, and *Bathyarchaeota* (Fig 6b). This community differed substantially from the communities in the SMTZ and MRZ, which were fairly similar to each other. In both sediment depths, *Bathyarchaeota*, *Chloroflexi*, and *Proteobacteria* were the dominant phyla.

## Discussion

### Identities of extracellular peptidases present in White Oak River Estuary sediments

Kinetics of fluorogenic substrate hydrolysis was consistent with the Michaelis-Menten rate law and hydrolysis rates were dramatically slower in autoclaved controls than in “live” treatments, indicating that the substrates were hydrolyzed by enzymes rather than by abiotic factors. The enzyme substrates used here encompassed a diverse range of amino acid and peptide chemistries, including polar and non-polar R groups at the P1 site (i.e., the amino acid N-terminal to the scissile bond) and substrates with and without steric protecting groups, which must have been hydrolyzed by endopeptidases (which cleave proteins from within) and aminopeptidases (which cleave proteins from the N terminus) respectively. Peptide bonds adjacent to a diverse set of amino acid residues were cleaved, including glycine (the smallest amino acid), phenylalanine (among the largest amino acids), arginine (positively charged at porewater pH) and leucine (uncharged, hydrophobic), consistent with the presence of a diverse range of extracellular peptidases throughout the core.

The metagenomic results also indicated the potential for a diverse range of secreted peptidases, produced by a broad range of taxa, throughout the sediment column (Fig 6). The metagenomic results represent a minimum estimate for the genomic potential for extracellular peptidase production, because they rely on the assumption that only those peptidases associated with signal peptides (SPs) are secreted. Non-SP-based enzyme secretion pathways may also contribute to the pool of extracellular enzymes, including Sec-independent secretion systems (30) and release of internal enzymes into the extracellular medium by viral lysis (31, 32).

The dominance of genes for exported gingipain-like endopeptidases (class C25) at all depths is consistent with rapid hydrolysis rates of fluorogenic substrates for endopeptidases. Gingipains are endopeptidases with preference for arginine at the P1 position (i.e., the N-terminal side of the hydrolyzed bond), which would be active towards the substrates GGR-AMC and Boc-VPR-AMC. Those were among the fastest-hydrolyzed substrates at each depth (Figs. 1, 2), indicating that genes for C25 peptidases were likely expressed. Previously, gingipains have been identified in Thermoprofundales (formerly Marine Benthic Group D) and in Bathyarchaeota, and appear to be widespread in marine sediments (17, 19, 33). The M28 family, also among the most abundant annotations, contains a diverse range of aminopeptidases and carboxypeptidases including leucine aminopeptidase, consistent with the observed hydrolysis of Leu-AMC (34). Genes for D-aminopeptidases genes were observed, consistent with hydrolysis of D-Phe-AMC.

Other abundant genes were annotated as methionine aminopeptidase, zinc carboxypeptidase, a C-terminal domain from bacterial pre-peptidases, and peptidases from the MEROPS families M24, S24, M50, and M28. Potential activities of these peptidases were not assayed. Zinc carboxypeptidases (M20) cleave enzymes from the carboxy-terminus and have strong specificity for Gly at the P1 position (i.e., the position C-terminal to the scissile bond), but little preference for the residue at the P1’ position (the position C-terminal to the scissile bond; in a carboxypeptidase this would be the C-terminus of the protein). Methionine aminopeptidases (M24) are metallopeptidases with preference for glycine at the P1 position.

S24 and M50 peptidases are less likely to be directly relevant to organic matter processing. S24 peptidases are involved in the SOS response for single-stranded DNA repair (34). M50 peptidases are membrane-bound enzymes that act as sporulation factors in Bacillus subtilis, and possibly other Bacteria (35, 36), and which are not secreted. However, DNA repair (37, 38) and spore formation (39, 40) both appear to be important survival mechanisms for microorganisms in subsurface sediments. The bacterial C-terminal pre-peptidase domain is often found in secreted peptidases, but is removed prior the peptidase becoming active, and could be associated with a wide range of peptidases (41).

Each of these annotations is plausible in terms of what is known about peptidase activities in sediments, and the annotations were generally consistent with the observed activities. We did not assay for carboxypeptidases (e.g. MEROPS family M20) or methionine aminopeptidase, but carboxypeptidases have previously been observed to be active in estuarine sediments (42) and the generally broad substrate specificities of extracellular aminopeptidases suggests that methionine aminopeptidases could have contributed to the hydrolysis of the other aminopeptidase substrates (43). However, a note of caution is also warranted when interpreting peptidase annotations from deeply branching microorganisms: the high diversity of hydrolases makes precise annotations difficult, and the exact substrate specificities of the peptidases in these samples may differ somewhat from those inferred from the annotations (44). Thus, while these annotations are environmentally plausible and generally consistent with the fluorogenic enzyme assays, they should nevertheless be viewed with some skepticism.

Interestingly, the sets of peptidases identified in genomes, and the activities observed, varied among depths much less than the microbes present (compare the similar profiles in Fig 6a to the notable differences among depths in 6b.) It is possible that extracellular enzymes were only produced by a small subset of taxa that are present at all sediment depths, although this would be inconsistent with previous evidence that diverse taxa including sulfate reducers and fermenters produce extracellular enzymes in sediments (albeit, deeper than those studied here, 19), the widespread phylogenetic distribution of similar extracellular enzymes (45), and previous observations of functional redundancy with respect to extracellular enzyme production in diverse systems (46–48).

### Peptidase kinetics suggest adaptaion of subsurface peptidases to degraded organic matter

Heterotrophic microorganisms in subsurface sediments have little access to fresh organic matter. In the cores described here, which represented ∼275 years of sediment deposition, organic matter oxidation rate decreased by at least three orders of magnitude between the surface and 82.5 cmbsf (Fig 5c). It is challenging to determine what fraction of high- vs low-molecular weight organic matter subsurface microorganisms metabolize. However, the fact that cell-specific *V_max_* was more-or-less constant downcore (Fig 5b) suggests that the heterotrophic community relied on complex organic matter to a similar degree at all depths. The cell-specific *V_max_* values for Leu-AMC hydrolysis, 21-51 amol cell^-1^ hr^-1^, are comparable to previous measurements in active environments such as surface sediments (2-100 amol cell^-1^ hr^-1^) and seawater (mostly less than 100 amol cell^-1^ hr^-1^, but with some measurements up to 10 nmol cell^-1^ hr^-1^; Vetter and Deming, 1994 and references therein), consistent with communities that relied primarily on organic carbon derived from macromolecules.

The ratio of Σ*V_max_* : OC oxidation rate is sensitive to the mix of enzymes included in the sum, and to the substrate specificity of enzymes assayed (some enzymes will be capable of hydrolyzing multiple substrates). The absolute value of that sum, therefore, is not particularly meaningful. The trend, however, is informative: as sediment depth increased, the potential activity of extracellular peptidases decreased much more slowly than the actual rate of organic carbon oxidation, so the ratio of Σ*V_max_* : OC oxidation rate increased dramatically (Fig 5d). *V_max_* is a proxy for enzyme concentration, so the observed increase in Σ*V_max_* : OC oxidation rate combined with the trend in cell-specific Σ*V_max_* suggests that deeper heterotrophic communities exhibited similar demand for detrital OM, but that those enzymes returned bioavailable hydrolysate at a much slower rate because substrate concentrations were lower. The White Oak River subsurface communities were similar to their surface counterparts in terms of reliance on extracellular enzymes for bioavailable organic carbon, although subsurface metabolisms were considerably slower. However, enzyme kinetics and potential activities of D-phenylalanine aminopeptidase, L-phenylalanine aminopeptidase, and L-ornithine aminopeptidase all suggested microbial community adaptation to old, degraded organic matter in deeper sediments.

Most amino acids are biosynthesized as L-stereoisomers. As organic matter ages, the ratio of D-amino acids to L-amino acids (D:L ratio) increases with depth, due to abiotic racemization and increased abundance of D-amino acids derived from bacterial cell walls (1, 50). Accordingly, the potential activity of D-phenylalanyl aminopeptidase increased relative to that of L-phenylalanyl aminopeptidase, indicating an increased capacity to access degraded organic matter. Ornithine, which is a product of the release of urea from arginine, is another marker for degraded organic matter, while phenylalanine is more characteristic of fresher organic matter (51), and the relative potential activity of L-phenylalanine aminopeptidase : ornithine aminopeptidase followed the same increasing trend with depth. Finally, the decrease of *K_m_* values with increasing depth indicates peptidases that function more efficiently at lower substrate concentrations. It is intuitive that the concentration of enzyme-labile organic matter concentrations would decrease downcore, and the observed increase in Σ*V_max_* : OC oxidation rate provides direct evidence of that. Taken together, these three observations provide strong evidence for a subsurface heterotrophic microbial community that is increasingly adapted to persist using degraded organic matter at increasing depth.

This evidence raises the ecological question of *how* selective pressure produces a heterotrophic community adapted to degraded organic matter. Modeling and genomic observations in older (thousands to millions of years), deeper sediments suggest that microbial growth rates are too slow for community adaptation by enhanced growth rates of more successful taxa; rather, communities in deeper sediments consist of taxa that were deposited at the sediment-water interface and died at the slowest rates (6, 52, 53).

If those findings can be generalized to the shallower environments investigated here, that poses a question: in which aquatic environments are microorganisms capable gaining reproductive advantage by growing on recalcitrant organic carbon? The studies cited above addressed sites at which sedimentation rates, microbial respiration, and likely cell doubling times were considerably slower than in the sediments described here, so even if growth (as opposed to persistence) on recalcitrant organic carbon is not possible in those environments, it may have been in the White Oak River sediments. Alternately, microbial taxa may gain adaptations to metabolize recalcitrant organic matter in environments where labile organic matter is more abundant and growth rates are higher. This scenario would imply that organisms which primarily metabolize more labile organic matter would gain some selective advantage by also metabolizing recalcitrant organic matter. Finally, it is not entirely clear how the microorganisms in this study used the amino acids resulting from extracellular hydrolysis. In deeper sediments, heterotrophs appear to be energy-limited rather than carbon limited (54), which would suggest that amino acids would be likely assimilated directly into proteins. However, if amino acids are catabolized, there may be an energetic advantage to incorporating D-amino acids: some catabolic pathways for L-amino acids involve conversion to the D-form prior to further processing, in which case uptake of D-amino acids could save energy (55). Further analysis of the mechanisms by which subsurface heterotrophs access degraded sedimentary organic matter may yield insights into how microorganisms survive in low-energy environments, and into the processes that shape the pool of organic carbon that is preserved or oxidized over geological timescales.

## Materials and Methods

### Study site

Samples were collected from Station H in the White Oak River Estuary, 34° 44.490’ N, 77° 07.44’ W, first described by Gruebel and Martens (56). The White Oak River Estuary occupies a drowned river valley in the coastal plain of North Carolina. Station H is characterized by salinity in the range of 10 to 28 and water depth on the order of 2 m (21). The flux of ΣCO_2_ across the sediment-water interface was 0.46 ± 0.02 mmol m^-2^ hr^-1^ (measured in May of 1987), primarily due to organic carbon oxidation via sulfate reduction. The sediment accumulation rate averages 0.3 cm yr^-1^. Total organic carbon content is approximately 5% (21). For this study, push cores of 40-85 cm were collected from Station H by swimmers on May 28, 2013, and October 22, 2014. In 2013, cores were transported to the nearby Institute of Marine Sciences (University of North Carolina) at Morehead City, where they were sectioned and processed for enzyme activities, porewater geochemistry, and cell counts within 6 hours of sample collection. Porewater sulfate in 2013 was depleted by 43.5 cm, and methane peaked at 79.5 cm (Fig. S1). In 2014, cores were transported on the day of sampling to the University of Tennessee, Knoxville, stored at 4 °C, and processed for enzyme activities the following day. Samples for metagenomic analysis were collected separately in October 2010 from three sites (sites 1, 2, and 3, as previously described by Baker et al., 2015), all of which are within 550 m of Station H. Porewater geochemistry of those samples is described in Figure S2 of Lazar et al (57). We note that, although the samples for sequencing were taken at a different time and slightly different location than the samples for enzyme assays, geochemistry and geomicrobiology of White Oak River sediments appears to be extremely stable and homogenous, possibly due to the fact that sediments are extremely fine and the estuarine water flow rates are low. In any case, geochemistry of the sediments are stable over timescales of decades (21, 24, 58) and microbial abundances in the SMTZ were very similar even though they were collected at sites separated by ∼500m (error bars in Fig 6b are mostly smaller than the differences among depths).

### Enzyme assays

Enzyme assays were performed using different protocols in 2013 (data presented in Figs 1-3) versus 2014 (data presented in Fig 4). In 2013, enzyme assays were performed according to a protocol similar to the one described in Lloyd et al (2013b). Cores were sectioned into 3 cm intervals. The following intervals were selected for enzyme assays: 0-3 cm, 3-6 cm, 27-30 cm, 57-60 cm, and 81-83 cm. Each section was homogenized, and approximately 0.5 ml wet sediment was transferred into separate 5 ml amber glass serum vials, which had been pre-weighed and preloaded with 4 ml anoxic artificial seawater (Sigma Sea Salts, salinity = 15, pH=7.5) Samples were weighed again to determine the precise mass of wet sediment added, and then an appropriate quantity of 20 mM peptidase substrate stock dissolved in DMSO was added, up to 90 μL, for final substrate concentrations of 0, 25, 50, 75, 100, 200, or 300 μM. Substrates are listed in Table 1. Triplicate incubations with 400 μM Arg-AMC, Gly-AMC, Leu-AMC and Gly-Gly-Arg-AMC were also created, but these were omitted for Ala-Ala-Phe-AMC and Boc-Phe-Val-Arg-AMC because the latter two substrates are considerably more expensive. Each serum vial was vortexed, briefly gassed with N_2_ to remove oxygen introduced with the sample, and approximately 1.3 ml slurry was immediately removed, transferred to a microcentrifuge tube, and placed on ice to quench the reaction. The precise time of quenching was recorded. This was centrifuged at 10,000 × *g* within approximately 15 minutes. The supernatant was transferred to a methacrylate cuvette and fluorescence was measured with a Turner Biosystems TBS-380 fluorescence detector set to UV mode (λ_ex_=365-395 nm, λ_em_=465-485 nm). Samples were then incubated at 16 °C, approximately the *in situ* temperature, and the sampling procedure was repeated after approximately 3 hours. The rate of fluorescence production was calculated as the increase in fluorescence for each sample divided by the elapsed time between sample quenching. Killed controls were made using homogenized, autoclaved sediments from 35-45 cmbsf. However, we note that autoclaving does not completely destroy sediment enzymes because sorption to mineral surfaces stabilizes enzyme structure, vastly increasing their ability to maintain a functional conformation at high temperatures (59–61). We therefore used the autoclaved samples as a qualitative control for the null hypothesis that enzymes were responsible for none of the observed substrate hydrolysis, rather than as a quantitative method to distinguish enzymatic substrate hydrolysis from potential abiotic effects. In some sediments, a large fraction of fluorophore can sorb to particles, requiring a correction to observed fluorescence (15, 16). In order to test the extent of sorption, we incubated 120 nM AMC in WOR sediment slurry (12.5 g sediment / 100 mL ASW, pH = 7.5) over the course of 125 hours. Fluorescence was stable over the entire incubation (Fig S2). Further, we sometimes measured fluorescence calibration curves repeatedly over ∼3 hours, and observed no clear changes in the slopes of the curves. We thus concluded that sorption of the free fluorophore was negligible.

In 2014, enzymes were assayed using a protocol based on the approach of Bell et al (2013), which was designed for soil enzyme assays. In this approach, peptidase substrates were mixed with sediment-buffer slurries in 2-mL wells of a deep-well plate. These plates were periodically centrifuged and 250 μL aliquots of supernatant were transferred into a black 96-well microplate. Fluorescence was read using a BioTek Cytation 3 microplate reader (λ_ex_ = 360 nm, λ_em_ = 440 nm). Results from this method proved considerably noisier than the single-cuvette method used in 2013, so kinetic parameters (*V_max_* and *K_m_*) were not calculated for these data. Nevertheless, results were qualitatively similar to those from 2013, and we have reported *V_max_* from 2014 as v0 measured at 400 μM substrate concentration, which was saturating. In October 2014, the following substrates were assayed: AAF-AMC, Arg-AMC, Boc-VPR-AMC, D-Phe-AMC, Gly-AMC, Leu-AMC, L-Phe-AMC, Orn-AMC, Z-Phe-Arg-AMC, and Z-Phe-Val-Arg-AMC. In October 2014, L-Phe-AMC, D-Phe-AMC, and Orn-AMC were assayed according to the same protocol in 3-cm core sections at 1.5, 4.5, 7.5, 10.5, 19.5, 22.5, 25.5, 28.5, 34.5, 37.5, 40.5, 43.5, 49.5, 52.5, 58.5, and 61.5 cmbsf.

Peptidase kinetic data were analyzed using R. All raw data and scripts related to enzyme analysis are posted at http://github.com/adsteen/WOR_enz_2013_2014. For samples taken using the more sensitive single-cuvette method, Michaelis-Menten parameters were estimated from nonlinear least squares fits to kinetic data. In the case of Leu-AMC at 4.5 centimeters below seafloor, kinetic data could not successfully be fit to a Michaelis-Menten function, so no *K_m_* was reported and the value of *v_0_* at the highest substrate concentration was substituted for *V_max_*. For analysis of correlations, data sets were qualitatively evaluated for homoskedasticity and normality of residuals using q-q plots and plots of residuals vs fitted values. When untransformed data met those criteria, the null hypothesis of no correlation was tested using linear least-squares regressions. When untransformed data failed to meet those criteria, data were log-transformed. In cases in which log-transformed data were either heteroskedastic or residuals were non-normally distributed, data were rank-transformed and correlations were tested using Spearman’s *ρ*.

### Geochemical and microbiological measurements

Sediment porosity was measured by mass after drying at 80 °C, according to the equation

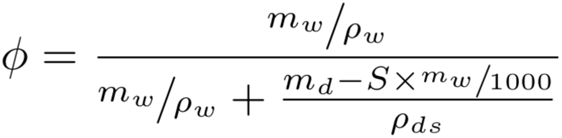

Here, *m_w_* represents mass lost after drying, *ρ_w_* represents the density of pure water, *m_d_* represents the mass of the dry sediment, *S* represents salinity in g kg^-1^, and *ρ_ds_* represents the density of dry sediment (assumed to be 2.5 g cm^-3^). Porewater sulfate concentrations were measured using a Dionex Ion Chromatograph (Sunnyvale, CA) in porewater that was separated by centrifugation in 15 ml centrifuge tubes at 5000 × *g* for 5 minutes, filtered at 0.2 μm, and acidified with 10% HCl. Methane was measured using 3 ml sediment subsamples that were collected from a cutoff syringe, entering through the side of a core section, immediately after core extrusion. Subsamples were deposited immediately in a 20 ml serum vial containing 1 ml, 0.1 M KOH. These were immediately stoppered and shaken to mix sediment with KOH. Methane was later measured by injecting 500 μl of bottle headspace into a GC-FID (Agilent, Santa Clara, CA) using a headspace equilibrium method (63).

### Geochemical modeling

Organic carbon remineralization rates as a function of depth were estimated by applying a multi-component reaction-transport model to depth distributions of sulfate and methane concentration. The model is based on equations described in Boudreau (64) and includes only sulfate reduction and methane production due to lack of data regarding oxic and suboxic processes. Thus, the model is limited to depths greater than 4.5 cm where sulfate reduction and methane production are the dominant processes, and bioirrigation and bioturbation may be assumed to be negligible. The organic matter remineralization rate is parameterized using the multi-*G* model first proposed by Jørgensen (65); a two-component model was sufficient to accurately simulate the sulfate and methane data. For solutes, the upper boundary conditions were measured values at 4.5 cm while the lower boundary conditions (200 cm) were set to zero-gradient. The flux of reactive organic carbon to 4.5 cm was calculated from the sulfate flux across the 4.5 cm horizon and an estimate of methane burial below the lower boundary (the methane flux at the upper boundary was observed to be zero), with an assumed oxidation state of reactive carbon of -0.7. The model contains four adjustable parameters that are set to capture the major details of measured sulfate and methane data: first-order rate constants for both fractions of the reactive carbon pool; the partitioning factor for both fractions, and the rate constant for methane oxidation.

### Cell enumeration

Cells were enumerated by direct microscopic counts. One mL of sediment was placed in a 2-mL screw-cap tube with 500 μl of 3% paraformaldehyde in phosphate buffered saline (PBS), in which it was incubated overnight before being centrifuged for 5 minutes at 3000 × *g*. The supernatant was removed and replaced with 500 μl of PBS, vortexed briefly and centrifuged again at 3000 × *g*. The supernatant was subsequently removed and replaced with a 1:1 PBS:ethanol solution. Sediments were then sonicated using a Branson Ultrasonics Sonifier SFX150 at 20% power for 40 seconds to disaggregates cells from sediments and diluted 40-fold into PBS prior to filtration onto a 0.2 μm polycarbonate filter (Fisher Scientific, Waltham, MA) and mounted onto a slide. Cells were stained with 4’,6-diamidino-2-phenylindole (DAPI) and enumerated by direct counts using a Leica Epifluorescence Microscope.

### Metagenomic analysis

To resolve the taxonomic distribution of extracellular peptidases we searched a pre-existing White Oak River *de novo* assembled and binned metagenomic dataset (Table S2; Baker *et al.*, 2015) for genes that were assigned extracellular peptidase functions. These assignments were based on best matches to extracellular peptidases in KEGG, pfam, and NCBI-nr (non-redundant) databases using the IMG annotation pipeline (Markowitz et al., 2014). Genes were additionally screened for signal peptidase motifs using the following programs: PrediSI setting the organism group to gram-negative bacteria (66), PRED-Signal trained on archaea (67), the standalone version of PSORT v.3.0 trained against archaea (68), and SignalP 4.1 using gram-negative bacteria as the organism group (69). All programs were used with default settings if not stated otherwise. Binned genomes from three different depth zones of White Oak River sediments were examined. The sulfate-rich zone (SRZ) genomes were obtained from sites 2 and 3 core sections 8-12 and 8-10 cm, respectively. The sulfate-methane transitions zone (SMTZ) genomes were recovered from site 2 and 3 and depths of 30-32 cm and 24-28 cm. The methane-rich zone (MRZ) was from site 1 and 52-54 cm. To determine abundance of phyla, we used the 16S rRNA gene sequences that were automatically extracted from the Baker et al metagenomes by IMG/m. We used CLC Genomic Workbench 10.0 (CLC Bio, Aarhus, Denmark) to trim adaptors and make contigs from bidirectional sequences. We clustered operational taxonomic units (OTUs) at 97% similarity used Silva reference set 132 to identify the taxonomy of OTUs (70).

## Acknowledgements

We thank Michael Piehler for access to his laboratory facilities at the University of North Carolina Institute of Marine Sciences, the captain of the R/V Capricorn for sampling assistance, and Terry Hazen for use of lab equipment at the University of Tennessee. We greatly appreciate two anonymous reviewers, whose constructive comments substantially improved the manuscript. Oliver Jeffers reminds ADS that science is cool and worth doing. Funding for KHM was provided by NSF grant DBI-1156644 to Steven W. Wilhelm. Funding for ADS was provided by NSF grant OCE-1431598 and a C-DEBI subaward. This work is C-DEBI Contribution number <<to be determined.>>

## Supplemental Information

Supplemental file: annotations of all peptidases included in this manuscript.

