## Supplementary material for "Kinetics and identities of extracellular peptidases in subsurface sediments of the White Oak River Estuary, NC": Figure S1 and S2

Supplemental figures.

Fig 1: Sulfate and methane profiles that were used to drive the model of OM remineralization rates presented in Fig 5.

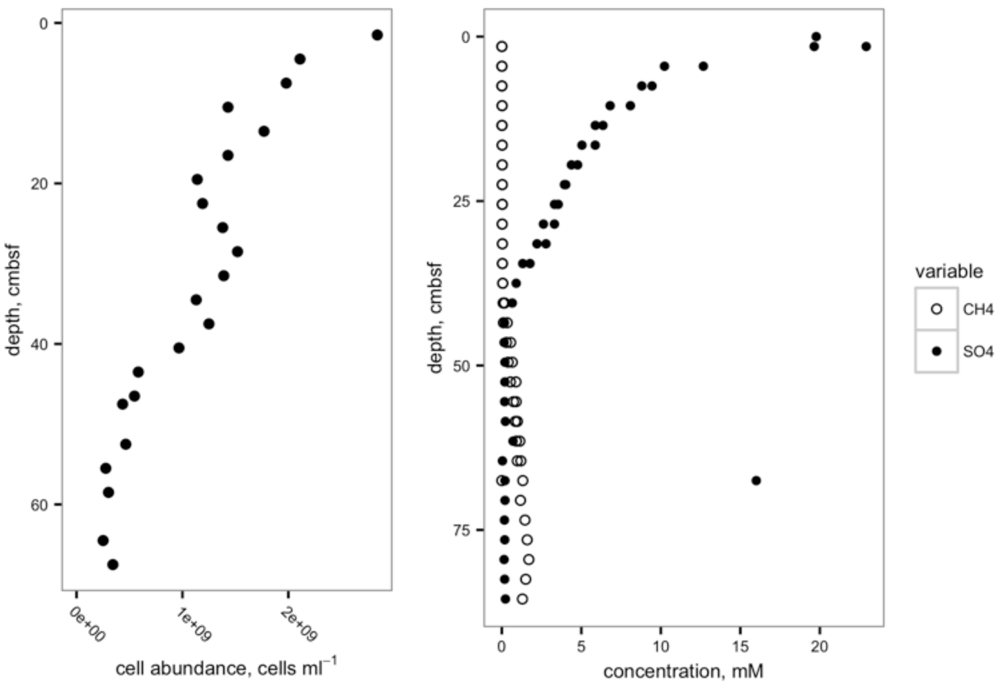

Fig 2: Sorption of 120 nM free AMC to White Oak River sediments in a slurry with ASW, salinity-18, pH 7.5. The value labeled as 0 incubation time is actually 120 nM AMC in ASW buffer without sediments. We attribute the 11% decrease in the presence of sediments to quenching by chromophoric OM, and interpret the stability of fluorescence in the presence of the slurry as evidence that no substantial sorption occurs. "rawFl" and "normFl" refer to data raw fluorescence data and data that are corrected for variability in the measured fluorescence of the solid standard respectively, which was negligible.

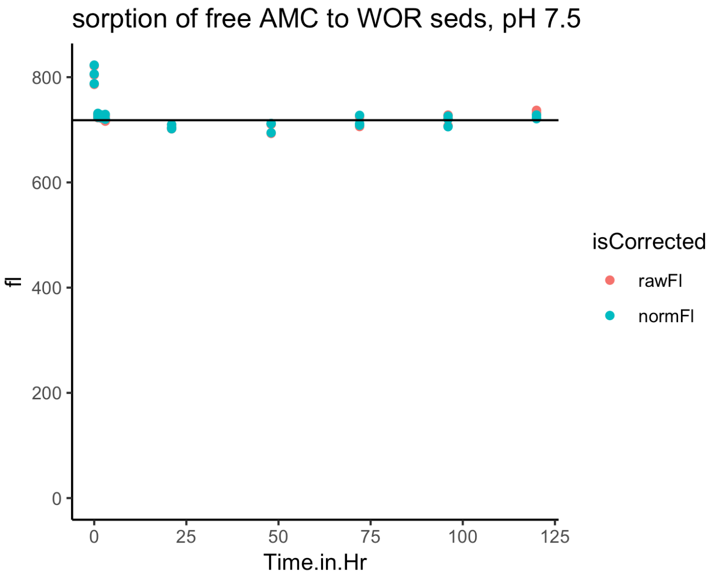
